# Robotic automation of droplet microfluidics

**DOI:** 10.1101/278556

**Authors:** Tuan M. Tran, Samuel C. Kim, Adam R. Abate

## Abstract

Droplet microfluidics enables new reactions, assays, and analytic capabilities, but often requires complex workflows involving numerous steps of macro- and micro-fluidic processing. We demonstrate robotically-automated droplet microfluidics, an approach to automate workflows with commercial fluid-handling robots. These workflows can be performed without human intervention, increasing reliability and convenience.

---

Droplet microfluidics enables new assay and analytic capabilities by performing reactions in compartmentalized emulsions, including accurate DNA and protein quantitation with digital droplet PCR (ddPCR) and ELISA (ddELISA), low-input DNA sequencing with digital droplet MDA (ddMDA), and ultrahigh-throughput single cell sequencing with droplet barcoding.^1–4^ These examples have been amenable to commercialization because they use simple workflows that only involve microfluidic droplet generation. However, applications such as cell phenotypic screening and nucleic acid cytometry use more complex workflows involving multiple macro- and micro-fluidic steps, including reagent preparation, cell encapsulation, pico-injection, and droplet sorting.^5–7^ Consequently, such workflows are limited to expert microfluidics labs. Moreover, other valuable workflows can be envisioned that are too complex even for expert labs to conduct.

To broaden the impact of droplet microfluidics, strategies for streamlining complex workflows are needed. One approach is to implement automated controllers consisting of pressure and temperature regulators, and integrated membrane valves;^8–11^ while this simplifies device operation, it remains difficult to introduce many reagents into numerous devices, as needed for most molecular biology applications. Alternatively, a custom “world-to-chip” interface consisting of a circular array of wells can programmatically infuse many reagents into a microfluidic device in a defined sequence.^12^ However, the instrument must be constructed from scratch and lacks flexibility for performing other important macroscopic fluid-handling processes, like combining, purifying, and analyzing reagents.

Indeed, robotic fluid-handling is a mature technology for automating numerous experiments in biology labs.^13–15^ Robotic instruments can execute complex workflows with precision and reproducibility exceeding that of a human. Moreover, they outclass custom microfluidic handlers in their capabilities and can interface with important laboratory hardware, like thermal incubators, reaction purifiers, and analysis tools, including optical, chromatographic, and mass spectrometry instruments.^16–19^ Currently, no microfluidic “lab on a chip” can match these instruments in capability and flexibility. Rather than reinventing automation, a superior approach would leverage existing robotics to enable both streamlining of droplet microfluidic workflows and new reaction and analysis tools.

In this paper, we present robotically-automated droplet microfluidics (RAD Microfluidics) an approach to automate droplet microfluidics using commercial fluid-handling robotics. Our approach consists of three components, the fluid-handling robot, a modular microfluidic system, and an interface for shuttling reagents between them. The system is controlled by a master computer to perform the requisite macro- and micro-fluidic operations for a given workflow. To illustrate the power of RAD Microfluidics, we use it to automate two workflows, ddPCR and *in vitro* directed evolution. Our results highlight a path forward for automating increasingly complex droplet microfluidic workflows.

## Results and discussion

The RAD Microfluidic system consists of separate robotic and microfluidic instruments that fluidically communicate through a modular pump and valve system (Fig. 1a). A master computer controls all components, commanding the robot as needed to process reagents and infuse them into the microfluidic devices. For fluorescence-activated droplet sorting (FADS), the computer uses a separate sorting instrument, specifying sorting gates to recover select droplets, which are then transported back to the robot for processing (Fig. 1b).

**Fig 1.**
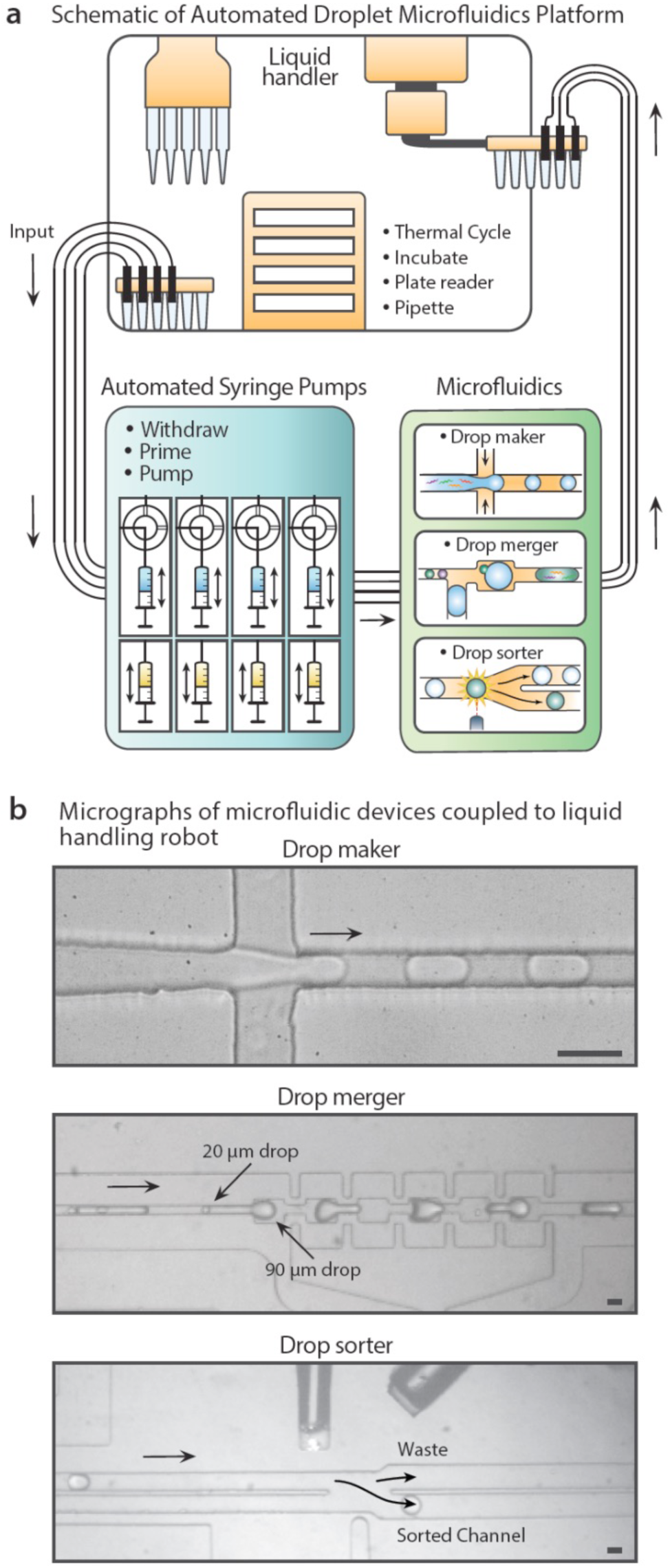
Overview of RAD microfluidics strategy. The instrument consists of three components, a commercial fluid-handling robot capable of processing, incubating, and analyzing fluids, a microfluidic breadboard consisting of common modules like droplet generators, picoinjectors, and sorters, and a fluidic communication highway consisting of arrays of pumps and valves (a). To use the microfluidic devices, the robot loads the requisite reagents into specialized wells connected to the pump array, aspirating and infusing them into the microfluidic devices at controlled flow rates (b). The entire instrument is controlled by a master computer. Scale bar is 50 μm.

As a demonstration of the approach, we use it to automate ddPCR, a simple workflow with high impact applications. The master computer instructs the robot to prepare and load ddPCR reagents and samples into pump reservoirs; hydrofluoroether (HFE) oil and surfactant for generating droplets are also loaded (Fig. 2a, left). The computer instructs the pumps to infuse the reagents into a microfluidic droplet generator, by reconfiguring rotary valves and flow in the pumps (Fig. 2a middle). Droplets are generated, traveling back to the robot through tubing, where they collect into a reservoir accessible to the robotic pipette. The robot exchanges HFE oil with thermostable FC40 and surfactant (Methods, Fig. 2a, right) and loads the emulsion into an onboard PCR machine for thermocycling (red box, Fig. 2a, right). This automated process can be repeated as desired, by programming the system to iterate on separate samples stored in well plates.

**Fig 2.**
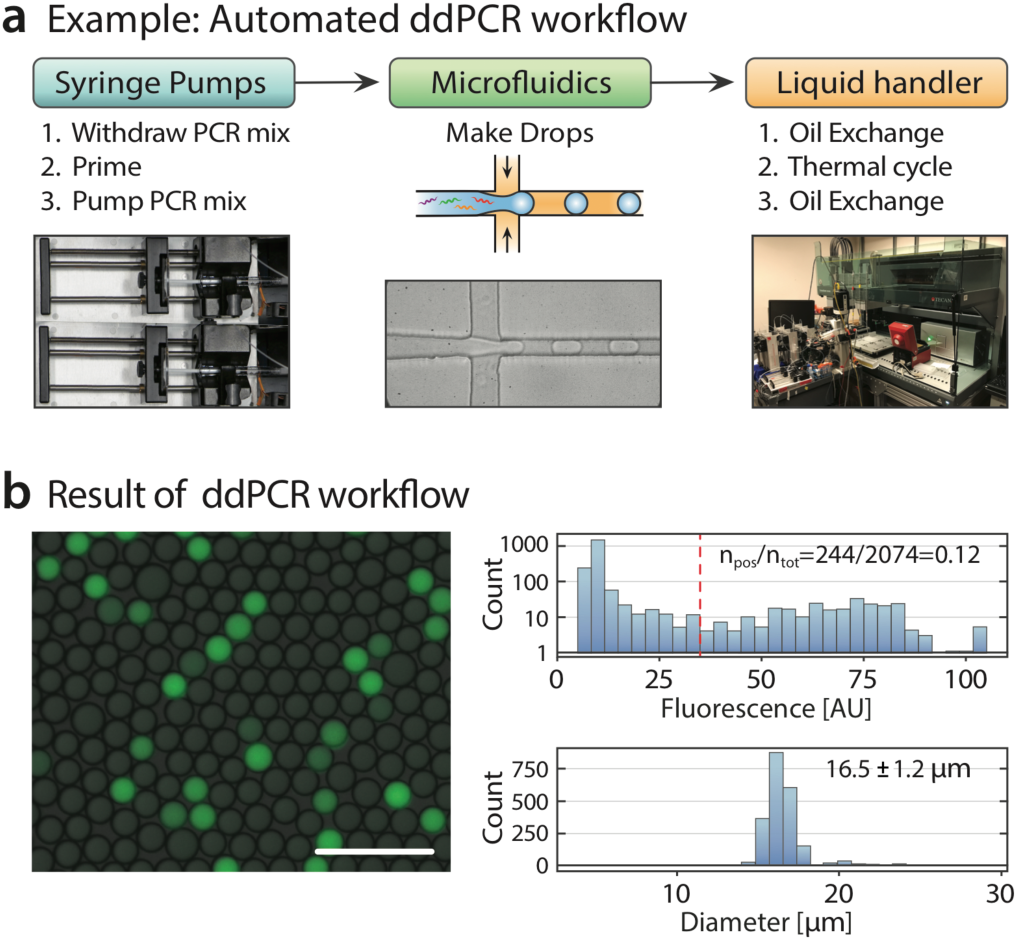
RAD Microfluidic automation of ddPCR. The ddPCR workflow consists of three steps, loading of robotically-prepared reagents into syringe pumps, encapsulation into monodisperse droplets via flow-focusing, and thermocycling of the emulsion aboard the robot (a). We image samples of the cycled droplets, observing that they are monodisperse and exhibit the characteristic “digital” fluorescence of ddPCR assays. Scale bar is 100 μm (b).

To assess amplification efficiency, we image samples of the droplets, observing the “digital” fluorescence characteristic of successful ddPCR (Fig. 2b, left and upper-right). The droplets are uniform in size, exhibiting polydispersity typical of thermocycled emulsions. The positive fraction (∼10%) agrees with the input DNA concentration, demonstrating that ddPCR is successful and can be used to infer target concentration. This shows that ddPCR can be automated with RAD Microfluidics, affording a strategy to scale the analysis of many samples.

To illustrate the flexibility of RAD microfluidics for complex workflows that are otherwise difficult to automate, we perform a mock *in vitro* evolution experiment using a workflow with droplet generation, merger, sorting, and multiple microfluidic handling operations. This workflow, and its variants, are valuable for applications of droplet microfluidics, including sequence-function mapping, *in vitro* enzyme evolution, and single-cell PCR, among others.^20–23^

The *in vitro* evolution process consists of iterative cycles of diversity generation (mutagenesis) and screening (droplet sorting). For mutagenesis, error-prone PCR can amplify a wild type sequence to generate a diverse library of mutants.^24,25^ Next, each mutant must be tested for activity, to identify enhancements. This can be accomplished by loading each mutant into a droplet containing a cell-free protein expression system (CFPE).^26–28^ The mutant sequence is *in vitro* translated to protein, generating a fluorescent signal to indicate enhancements when an appropriate assay is utilized. For example, this can be applied to enhance an enzyme, pathway, or genetic circuit.^29–31^ However, a single DNA molecule is usually insufficient for accurate mutant characterization. To enhance signal, mutant copy number in each droplet must be increased, which can be accomplished by programming the system to encapsulate and ddPCR amplify the error-prone library. The mutant sequences are loaded in droplets by flow-focusing under limiting dilution and amplified on the onboard PCR machine. We demonstrate the capability to automate the ddPCR process in Fig 2. For the mock experiment, we made FITC droplets of a similar size as a proxy for ddPCR droplets. (Methods, Fig. 3a). The resultant droplets are monodispersed (Fig. 3b) and exhibit the expected fluorescence (Fig. 3c).

**Fig 3.**
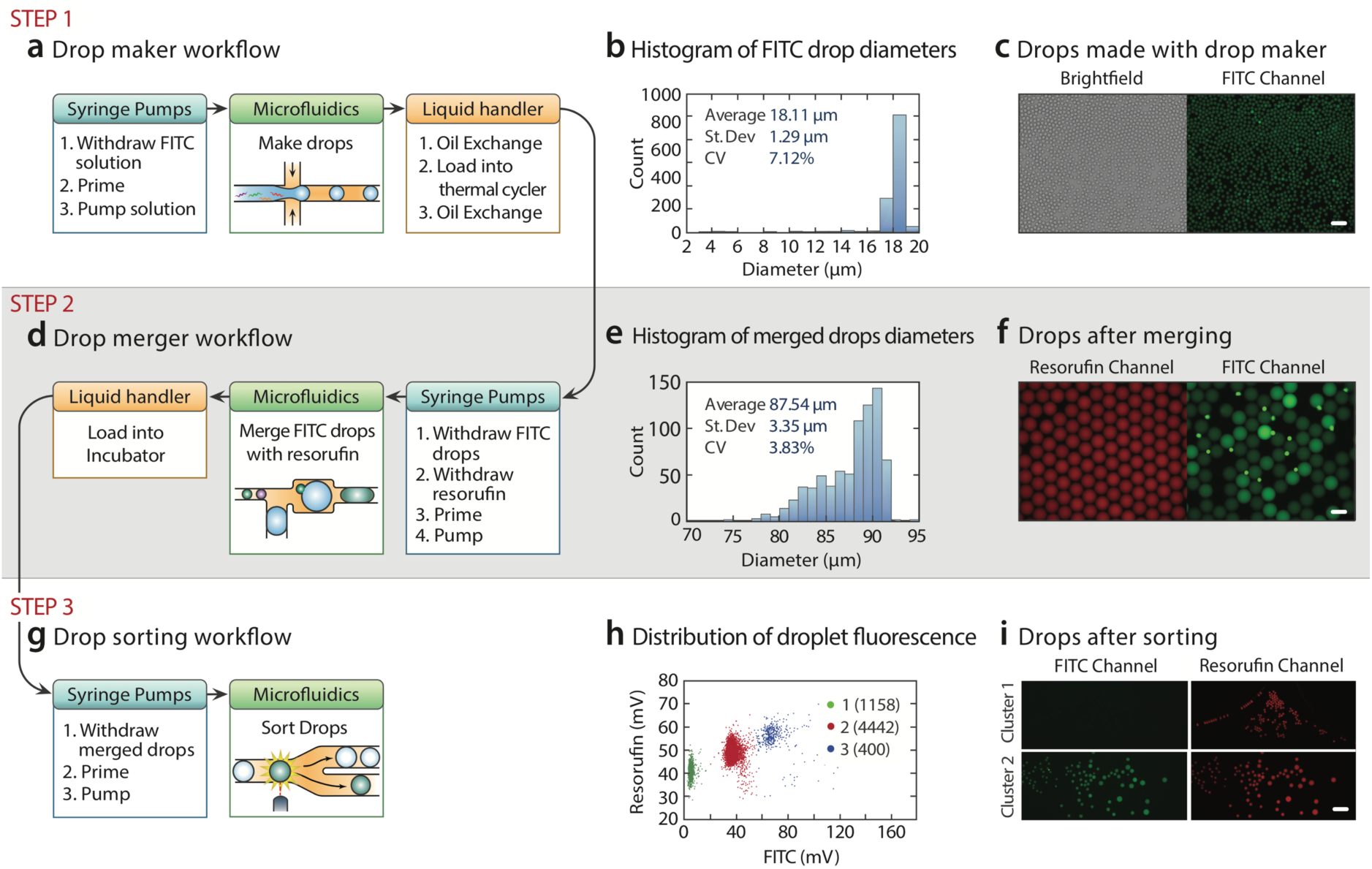
RAD Automation of multistep workflow used for *in vitro* evolution. DNA sample is encapsulated by flow focusing (a) and the monodispersed droplets (b) thermocycled with a PCR machine on the robot to digitally amplify each molecule, providing sufficient DNA for cell-free protein expression. Scale bar is 100 μm. (c). The amplified droplets are merged with CFPE droplets via an electrocoalescence device (d), exhibiting polydispersity characteristic of this imperfect process (e), although in fluorescence mode a large fraction of droplets appear to have properly paired and merged. Scale bar is 100 μm. (f). The merged droplets are incubated, to allow expression of green-fluorescent reporter, and then sorted via FADS (g). The presorted droplets exhibit three populations, unmerged, 1:1 merged, and multiply-merged, when viewed on a two-color fluorescence scatter plot (h). By gating specific populations, the instrument recovers droplets with the desired fluorescence properties. Scale bar is 200 μm. (i).

To assay each mutant for activity, CFPE media must be added to each droplet. As the second step in our workflow, we thus instruct the robot to perform pair-wise droplet merger (Fig. 3d). Dye-labeled CFPE droplets are generated in a T-junction, paired with reinjected FITC droplets, and coalesced.^32,33^ The merged droplets exhibit polydispersity characteristic of electrocoalescence, arising from instability of the emulsion upon collection and merger of too many or few droplets in the electrode region (Fig. 3e). Consequently, while most merged droplets are FITC-positive, indicating successful merger, intensity varies across the population, which includes small, unmerged FITC droplets (Fig. 3f, right). The droplets can be incubated at this stage if necessary, for instance, to allow protein translation and the assay to occur.

The final step in *in vitro* evolution is to sort the incubated droplets by fluorescence, to recover the most active variants, which our robot accomplishes via FADS. The droplets are injected into the sorter, quantified for size and fluorescence using a fiber-optic detector, and sorted to prescribed gates (Fig. 3g). The droplets exhibit three major populations: Cluster 1 with low FITC corresponding to cell-free extract droplets that either merged with no or negative ddPCR droplets; Cluster 2 with moderate FITC signal corresponding to desired 1:1 mergers; and Cluster 3 corresponding to droplets that have undergone multiple mergers (Fig. 3h). We instruct the system to sort Clusters 1 and 2 into separate wells, imaging the results (Fig. 3i). We find that droplets sorted for Cluster 1 are negative for FITC but positive for resorufin, while ones for Cluster 2 are double-positive, as expected based on the gating, demonstrating successful sorting.

These results show that RAD microfluidics can perform with automation all steps for an *in vitro* evolution workflow, including library generation, single mutant testing, and high-throughput sorting.

## Conclusions

A barrier to advancing droplet microfluidics is that workflows are often complex, requiring an expert to design and run them. RAD microfluidics automates workflow operation for increased ease and reliability. Moreover, by combining microfluidic components with robotic fluid-handling, the approach makes available new reaction and analytic capabilities difficult to integrate into existing droplet microfluidic devices, like sample purification, thermal cycling, and analysis.

The main challenge to implementing RAD microfluidics is the custom pump and valve system that interfaces between the robot and microfluidic devices. In addition, our simple prototype lacks fault handling, and thus error detection. Nevertheless, even in its nascent form, RAD microfluidics illustrates the value of removing humans from the execution of microfluidic workflows, allowing more reliable performance, and ultimately enabling increasingly complex workflows involving many macro- and micro-fluidic processing steps.

## Materials and methods

### Device fabrication

Microfluidic devices were fabricated using soft lithography. SU-8 3025 photoresist (MicroChem) was spincoated on a 3-inch silicon wafer, exposed and developed to make a master mold structure. Droplet maker, merger and sorter were fabricated to be 20, 60 and 90 μm tall. PDMS elastomer (RTV615, Momentive) was mixed at 10:1 ratio, poured on the master mold and cured at 65°C for 1 hour. PDMS slab was removed from the wafer by cutting and the access holes (0.75 mm in diameter) were punched. The channel side of the PDMS slab was plasma-bonded to a glass slide (12-550C, Fisher Scientific) by treating with oxygen plasma for 60 s at 1 mbar (PDC-001, Harrick Plasma). The inner surface of the microchannels was transformed to hydrophobic by flowing in Aquapel^®^.

### Digital droplet PCR

2% HFE oil is composed of HFE (Novec 7500, 3M) supplemented with 2%(w/w) 008-FluoroSurfactant (RAN Biotechnologies). 5% FC40 oil is FC40 (Sigma) supplemented with 5%(w/w) 008-FluoroSurfactant. 2% HFE oil was used to make a water-in-oil emulsion containing Phusion PCR master mix (M0530S, Thermo Fisher) and 70 fM template DNA (mCherry gene from plasmid pfm301; sequence shown in Supplementary Information) in the detergent-free buffer (F520L, Thermo Fisher) supplemented with 2.5%(v/v) Tween 20 and 2.5%(v/v) PEG 6000. Flow rates of aqueous and oil phases used for generating droplets were 125 and 250 μL/hr, respectively. The oil phase of collected droplets was exchanged with 5% FC40 oil by instructing the liquid handling robot to remove oil from the bottom of PCR tubes because the aqueous droplets float on top of the denser fluorinated oil. 5% FC40 oil supports better droplet stability over the course of the PCR thermal cycle. After PCR, the oil phase was exchanged again with 2% HFE oil supplemented with SYBR Green dye (S7563, Thermo Fisher) to stain the PCR products. The droplets were visualized on a microscope (EVOS FL, Thermo Fisher) using both transmission and GFP channels. The acquired images were analyzed with the ImageJ software to extract droplet size and fluorescence distributions.

### Multi-step workflow

1 μM solution of FITC-labeled dextran (D1845, Thermo Fisher) was automatically loaded into a syringe and emulsified using the same protocol as ddPCR. Then, the FITC droplets were collected into a 96-well plate by the liquid handler. The robot exchanged the oil phase of FITC drops from 2% HFE to 5% FC40. The robotic arm transferred the well plate to the thermocycler, which closed and performed a dummy cycle (no heating or cycling). After the cycle, the thermocycler opened, and the robotic arm moved the well plate back to the pipettor region. The liquid handler performed another oil exchange to revert back to 2% HFE and moved the well plate to automated syringes.

### Master scheduler design

Automation of droplet microfluidics requires automation of the syringe pumps and liquid handling. An illustration of a modular unit composed of a liquid handler, a pump/valve array and a microfluidic device is shown in Figure S1. The picture of the entire setup is shown in Figure S2. A web link to the video recording of the operational steps is shown in Movie S1 in the Supplementary Information.

1. A master scheduler was written in LabVIEW (National Instruments) to control when syringe pump or liquid handling operations are performed.
2. The master scheduler follows a list of steps set by the user.
  a. If it is a syringe pump operation, the scheduler can directly control the syringe pumps to carry out these operations.
  b. If it is a liquid handling operation, the scheduler simply instructs the liquid handler to perform a specific protocol pre-written in Tecan’s EVOware software.
  c. Coordination between the syringe pump operation and liquid handling is performed by creating text files. A text file called “run_fluidics.txt” is created automatically when Tecan is finished with its specific protocol. The labview scheduler will detect when “run_fluidics.txt” is created and proceed to run the next syringe pump operation. When the syringe pump operation is completed, a text filed called “complete.txt” is created. The creation of this file instructs Tecan to move on to its next pre-written protocol.

### Automated syringe pump system

Syringe pumps (NE-501, New Era Pump Systems) were connected to rotary valves from Tecan Cavro pumps (PN 20738707). The Tecan Cavro is capable of pumping but not at the slow flow rates required for droplet microfluidics. The syringe pumps and rotary valves were controlled through a custom LabVIEW software. The software rotates the valves to the appropriate position for aspiration or dispensing. It also controls the flow rate and volume dispensed by the syringe pumps.

### Liquid handler system

The liquid handler system (Freedom EVO, Tecan) was controlled through the software provided by the manufacturer (Freedom EVOware, build 2.6.17). Protocols for pipetting and liquid transfer are written in the Freedom EVOware software. After each fluidic operation, the robot communication with LabVIEW is achieved by monitoring text files in a shared network folder.

### Optical configuration

A fiber-optic setup was used for detecting fluorescence signals from droplets as previously described.^34^ This optical configuration allowed us to decouple the fluorescence measurement from the microscope. Unlike the conventional epi-fluorescence setup where both excitation and emission light paths share the same objective lens and the photodetector is often attached to the microscope, the fiber-based setup can have the light source and the detector that are physically separate from the microscope. This configuration is useful for monitoring droplet quality over the course of an experiment because the microscope can be used to visualize any part of the operated device while the fluorescence signal is acquired continuously from the separate fiber detection point.

## Acknowledgments

This work was supported by the Chan Zuckerberg Biohub, the National Science Foundation CAREER Award (Grant Number DBI-1253293); the National Institutes of Health (NIH) (Grant Numbers R01-EB019453-01, R01-HG008978, HG007233-01, DP2-AR068129-01, 1R21HG007233, and R21AI116218); and the Defense Advanced Research Projects Agency Living Foundries Program (Contract Nos. HR0011-12-C-0065, N66001-12-C-4211, HR0011-12-C-0066) and Fold F(x) Program (Contract No. DE-AC02-05CH11231). Work at the Molecular Foundry was supported by the Office of Science, Office of Basic Energy Sciences, of the U.S. Department of Energy under Contract No. DE-AC02-05CH11231.

